# Priming enables a NEK7-independent route of NLRP3 activation

**DOI:** 10.1101/799320

**Authors:** Niklas A. Schmacke, Moritz M. Gaidt, Inga Szymanska, Fionan O’Duill, Che A. Stafford, Dhruv Chauhan, Adrian L. Fröhlich, Dennis Nagl, Francesca Pinci, Jonathan L. Schmid-Burgk, Veit Hornung

**Affiliations:** Gene Center and Department of Biochemistry, Ludwig-Maximilians-Universität München, Munich, Germany; Center for Integrated Protein Science (CiPSM), Ludwig-Maximilians-Universität München, Munich, Germany; Department of Molecular & Cell Biology, and Cancer Research Laboratory, University of California, Berkeley, CA, USA; Broad Institute of MIT and Harvard, Cambridge, Massachusetts, USA

## Abstract

The NLRP3 inflammasome plays a central role in antimicrobial defense, as well as in sterile inflammatory conditions. NLRP3 activity is governed by two independent signals. The first signal primes NLRP3, allowing it to respond to its activation signal. In the murine system, the mitotic spindle kinase NEK7 has been identified as a crucial factor in relaying the activation signal to NLRP3. Here we show that the requirement for NEK7 can be bypassed by TAK1-dependent post-translational priming. Under pro-inflammatory conditions that activate TAK1, NEK7 was dispensable for NLRP3 inflammasome formation in human and murine cells. Intriguingly, dissecting the NEK7 requirement in iPSC-derived primary human macrophages revealed that this NEK7-independent mechanism constitutes the predominant NLRP3 priming pathway in these cells. In summary, our results suggest that NEK7 functions as an NLRP3 priming – rather than activation – factor that can work in synergy or redundancy with other priming pathways to accelerate inflammasome activation.

## INTRODUCTION

Cells of the innate immune system employ a repertoire of so-called pattern recognition receptors (PRRs) to discriminate self from non-self. Engagement of these PRRs triggers a broad array of effector functions that are geared towards eliminating a microbial threat. The inflammasome pathway constitutes a special class of this PRR system that is signified by the activation of the cysteine protease caspase-1 in a large supramolecular protein complex (Broz and Dixit, 2016). Activation of caspase-1 causes maturation of pro-inflammatory cytokines, most prominently IL-1β (Dinarello, 2018), as well as the induction of a special type of cell death, known as pyroptosis (Vande Walle and Lamkanfi, 2016). Among several inflammasome sensors, NLRP3 plays a pivotal role in antimicrobial defense as well as sterile inflammatory diseases (Patel et al., 2017). This is owed to the fact that NLRP3 is a highly sensitive, yet non-specific PRR. In this regard, NLRP3 has been shown to respond to the perturbation of cellular homeostasis by a broad array of diverse stimuli, rather than being activated by a specific microbe-derived molecule (Swanson et al., 2019). K^+^ efflux from the cytosol has been identified as a common denominator of many NLRP3 triggers (Muñoz-Planillo et al., 2013). In this function several types of lytic cell death have been shown to result in secondary engagement of the NLRP3 inflammasome pathway (Gaidt and Hornung, 2018). However, K^+^-efflux independent NLRP3 stimuli have also been described (Gaidt et al., 2016; Groß et al., 2016) and a recent report has identified dispersal of the trans-Golgi network (TGN) to be a common denominator of both potassium dependent- and independent NLRP3 triggers (Chen and Chen, 2018).

Unlike other inflammasome sensors, NLRP3 critically depends on the engagement of a priming step (Hornung and Latz, 2010). This priming signal can be provided by different types of receptors, typically PRRs that are able to trigger NF-kB activation. Lipopolysaccharide (LPS) activating TLR4 is commonly used to provide a priming signal preceding the actual NLRP3 activation step. Initially, the necessity of priming had been ascribed to the fact that NLRP3 is expressed at limiting levels in murine macrophages. In this respect it has been shown that NF-kB activating stimuli drive the expression of *Nlrp3*, thereby facilitating its activation (Bauernfeind et al., 2009; Kahlenberg et al., 2005). In line with these findings, inhibition of transcription blocks this mode of NLRP3 priming, whereas transgenic expression of NLRP3 bypasses the requirement of transcriptional priming (Bauernfeind et al., 2009; Kahlenberg et al., 2005). Extending this concept, NLRP3 can also be primed non-transcriptionally, e.g. by a short pulse of LPS treatment (Bauernfeind et al., 2016; Juliana et al., 2012; Lin et al., 2014). These modes of priming have been shown to depend on a variety of posttranslational modifications of NLRP3, including phosphorylation, de-phosphorylation, de-ubiquitination, and de-sumoylation (Barry et al., 2018; Shim and Lee, 2018). Although being mechanistically unrelated, these events are commonly referred to as post-translational priming and indicate that many cells already express sufficient amounts of NLRP3 to activate the inflammasome under steady-state conditions.

Despite considerable insight into pathways that mediate NLRP3 priming, the activation step of the NLRP3 inflammasome remained uncharacterized. In this regard, three reports have recently identified the mitotic spindle kinase NEK7 as a critical cofactor in NLRP3 activation in murine cells (He et al., 2016b; Schmid-Burgk et al., 2015; Shi et al., 2016). Notably, this role of NEK7 is distinct from its function in the cell cycle, as its kinase activity is not required for NLRP3 activation (He et al., 2016b; Shi et al., 2016). NEK7 has been suggested to interact with NLRP3 in a K^+^-efflux dependent manner, and deletion of NEK7 does not affect transcriptional NLRP3 priming (He et al., 2016b; Shi et al., 2016). This has led to the conclusion that NEK7 is involved in NLRP3 activation downstream of K^+^-efflux (He et al., 2016a). Of note, studies identifying NEK7 as an indispensable factor for NLRP3 activation have mainly been conducted in murine models. Here, we report that reductionist genetic dissection of NLRP3 signaling in human cells revealed a new pathway of NLRP3 activation that depends on post-translational modification and enables NLRP3 inflammasome activation independently of NEK7.

## RESULTS

### Human myeloid cell lines activate NLRP3 in the absence of NEK7

To investigate whether human cells also required NEK7 for NLRP3 activation we made use of the BLaER1 transdifferentiation system that we have previously adopted to study innate immune sensing (Gaidt et al., 2018; Rapino et al., 2013). We subjected *NEK7*^-/-^ BLaER1 cells to four hours of LPS priming and subsequently stimulated these cells with Nigericin (NLRP3) or Needle Tox (NAIP/NLRC4). As expected, NEK7-deficiency showed no impact on NAIP/NLRC4 inflammasome activation, as assessed by IL-1β maturation (Fig. 1A) or LDH release (Fig. 1B). Surprisingly, *NEK7*^-/-^ cells also showed no impairment of their NLRP3 inflammasome response (Fig. 1A, B). NEK6 is a close homologue of NEK7, sharing more than 80 % sequence identity. To address whether NEK6 was functionally redundant with NEK7 for NLRP3 activation in human cells, we generated cells deficient for both NEK6 and NEK7. Similar to NEK7-deficient cells, *NEK6*^-/-^, *NEK7*^-/-^ BLaER1 cells displayed unimpaired activation of the NLRP3 inflammasome (Fig. 1A, B). As expected, *NLRP3*^-/-^ cells showed no response to Nigericin stimulation, while they remained responsive to NAIP/NLRC4 inflammasome activation. In line with these observations, caspase-1 maturation upon Nigericin treatment also proceeded independently of NEK7 (Fig. 1C). To investigate if Nigericin stimulation still activated the NLRP3 inflammasome via potassium efflux in absence of NEK7, we firstly utilized the NLRP3-specific inhibitor MCC950 (Coll et al., 2015). Pretreatment with MCC950 blunted inflammasome activation in wildtype, *NEK7*^-/-^ and *NEK6*^-/-^, *NEK7*^-/-^ cells stimulated with Nigericin, while it left the NAIP/NLRC4 inflammasome intact (Fig. S1A-C). Secondly, preventing K^+^ efflux by increased extracellular K^+^ concentration was still able to block NLRP3 but not NAIP/NLRC4 signaling in absence of NEK7 (Fig. 1D and S1D), indicating that Nigericin still relied on inducing potassium efflux to trigger NLRP3 inflammasome activation. Finally, we sought to validate the observation of NEK7-independent NLRP3 activation in a different cell culture model and thus turned to THP-1 cells. Analogous to the results obtained in BLaER1 cells, THP-1 cells deficient in NEK7 showed no attenuation of Nigericin-triggered inflammasome activation, whereas *NLRP3*^-/-^ THP-1 cells were completely defective (Fig. 1E-G). Altogether, these results indicated that NEK7 and also NEK6 are not required for NLRP3 activation in human myeloid cell lines.

**Figure 1.**
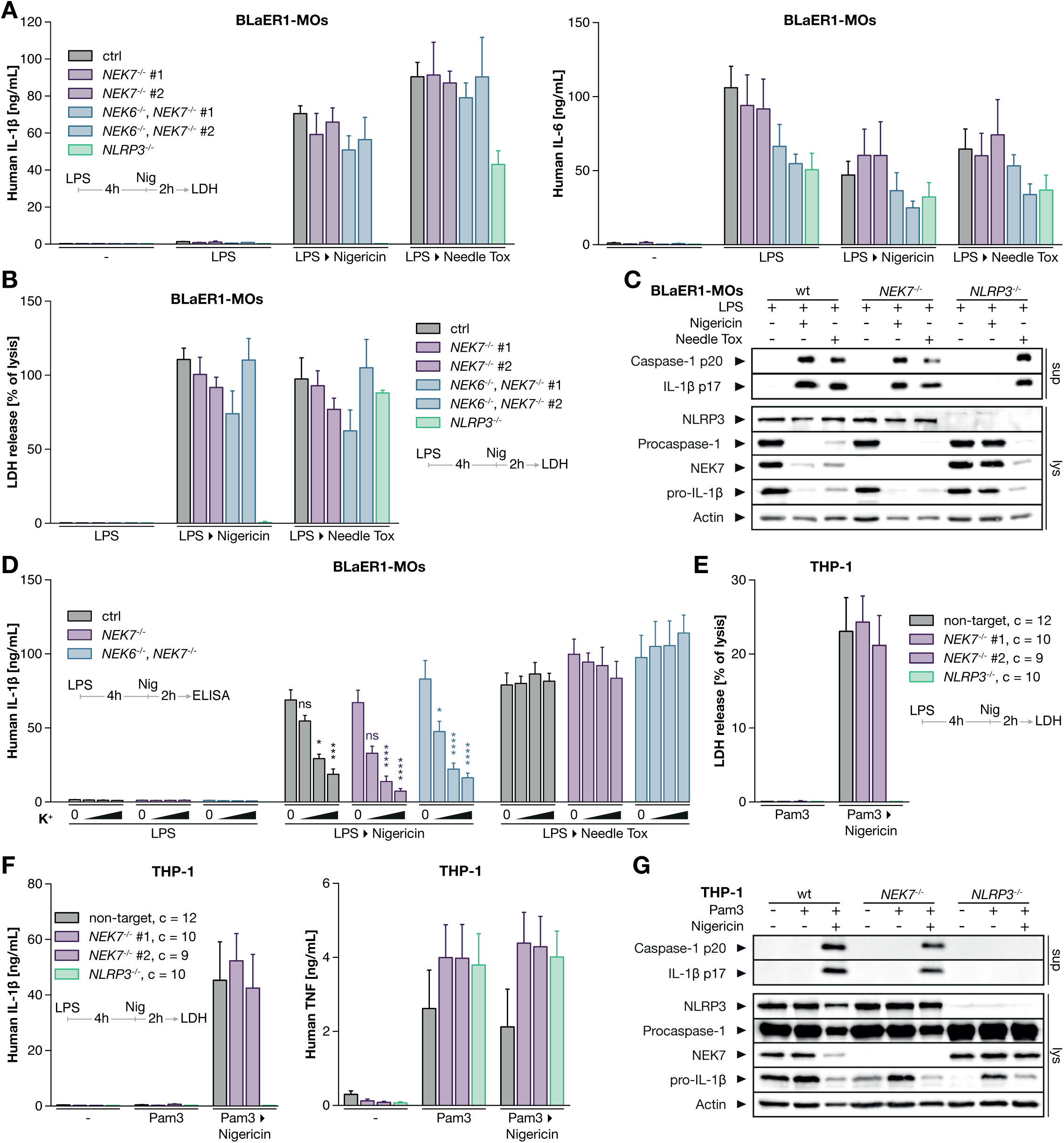
Human myeloid cell lines activate NLRP3 independently of NEK7. (A, B, C) BLaER1 monocytes of the indicated genotypes were primed with LPS for 4 hr and subsequently stimulated with Nigericin or pA and Needle Tox (LF-YscF). IL-1β, IL-6 (A) and LDH release (B) of two clones per genotype are depicted as mean + SEM of three experiments. (C) One representative immunoblot of three independent experiments is shown for caspase-1 release. (D) BLaER1 monocytes of the indicated genotypes were stimulated as in (A) in the presence of up to 60 mM potassium chloride. LDH release is depicted as mean + SEM of three experiments. (E, F, G) THP-1 cells of the indicated genotypes were primed with Pam3CSK4 for 4 hr and subsequently stimulated as in (A). Data are represented as mean + SEM of the indicated number of c=clones from one representative experiment of two. Two different sgRNAs targeting NEK7 are shown. The immunoblot in (G) is one representative of three independent experiments. **** p < 0.0001, *** p < 0.001, * p < 0.05, ns p ≥ 0.05. See also Figure S1.

### TAK1-priming bypasses NEK7 during NLRP3 inflammasome activation

The observation that human BLaER1 and THP-1 cells can activate NLRP3 independently of NEK7 suggests that another signaling event can compensate for NEK7. We wondered whether this compensatory signaling entity also existed in murine cells. To circumvent any effect of transcriptional NLRP3 priming, we addressed this question in murine macrophages that constitutively express NLRP3 (Franklin et al., 2014). As previously shown (Schmid-Burgk et al., 2015), Nigericin-mediated inflammasome stimulation did not require TLR-priming and was completely dependent on NEK7 under these conditions (Fig. 2A-C). However, concomitant stimulation with LPS enabled NEK7-independent NLRP3 inflammasome activation in these cells, suggesting that LPS-priming can also bypass the requirement of NEK7 in murine cells (Fig. 2A-C). Analogous results were obtained with R848 or Pam3CSK4 instead of LPS (Fig. S2A-D), and when ATP was used to activate the NLRP3 inflammasome (Fig. 2B). Further, NEK7-independent inflammasome activation in mouse cells was confirmed to drive caspase-1 maturation by immunoblot (Fig. 2C). Of note, simultaneous LPS treatment also enhanced NEK7-dependent NLRP3 inflammasome activation, especially when assessing short stimulation periods (Fig. S2E, F). In light of the constitutive Nlrp3 expression that renders these cells insensitive to transcriptional priming of NLRP3 these results suggest that LPS primes NLRP3 post-translationally to allow NEK7-independent activation. In line with this notion, the NEK7-bypass effect was not affected by treatment with the translation-inhibitor cycloheximide (Fig. 2D, S2G). To elucidate the signaling cascade of NEK7-independent post-translational priming, we genetically perturbed TLR4 and its downstream signaling adaptors TRIF (*Ticam1*) and Myd88 on either wt or *Nek7*^-/-^ backgrounds in these cells. *Ticam1*^-/-^ and *Myd88*^-/-^ cells displayed a selective lack of antiviral (IP-10) and pro-inflammatory (TNF) gene expression, whereas TLR4 deficiency abrogated LPS-dependent cytokine production altogether (Fig. 2E). LPS-dependent post-translational NLRP3-priming was dependent on TLR4, yet redundantly employed MyD88 and TRIF. Consequently, only the deletion of both MyD88 and TRIF phenocopied the TLR4 knockout and fully abrogated NEK7-independent NLRP3 activation (Fig. 2E). Since both these pathways converge at the level of the TAK1 complex, we next sought to investigate the role of TAK1 (*Map3k7*) activity. Since *Map3k7*^-/-^ murine macrophages were not viable, we instead turned to the small molecule inhibitor Takinib. As expected, macrophages treated with Takinib displayed markedly, but not completely, reduced TNF responses upon LPS treatment (Fig. 2F). Stimulating these cells with Nigericin alone only showed a NLRP3-dependent inflammasome response that was completely NEK7-dependent, while inhibiting TAK1 exerted no effect on this response. Conversely, stimulating these cells with LPS + Nigericin uncovered NEK7-independent NLRP3 inflammasome activation, whereas TAK1 inhibition blocked this effect to the same extent it blocked TNF production (Fig. 2F). Taken together, these data establish that post-translational LPS-priming employs TAK1 to alleviate the requirement for NEK7 in NLRP3 inflammasome activation. Moreover, these findings also indicate that TAK1 kinase activity is required for NEK7 independent PTM priming.

**Figure 2.**
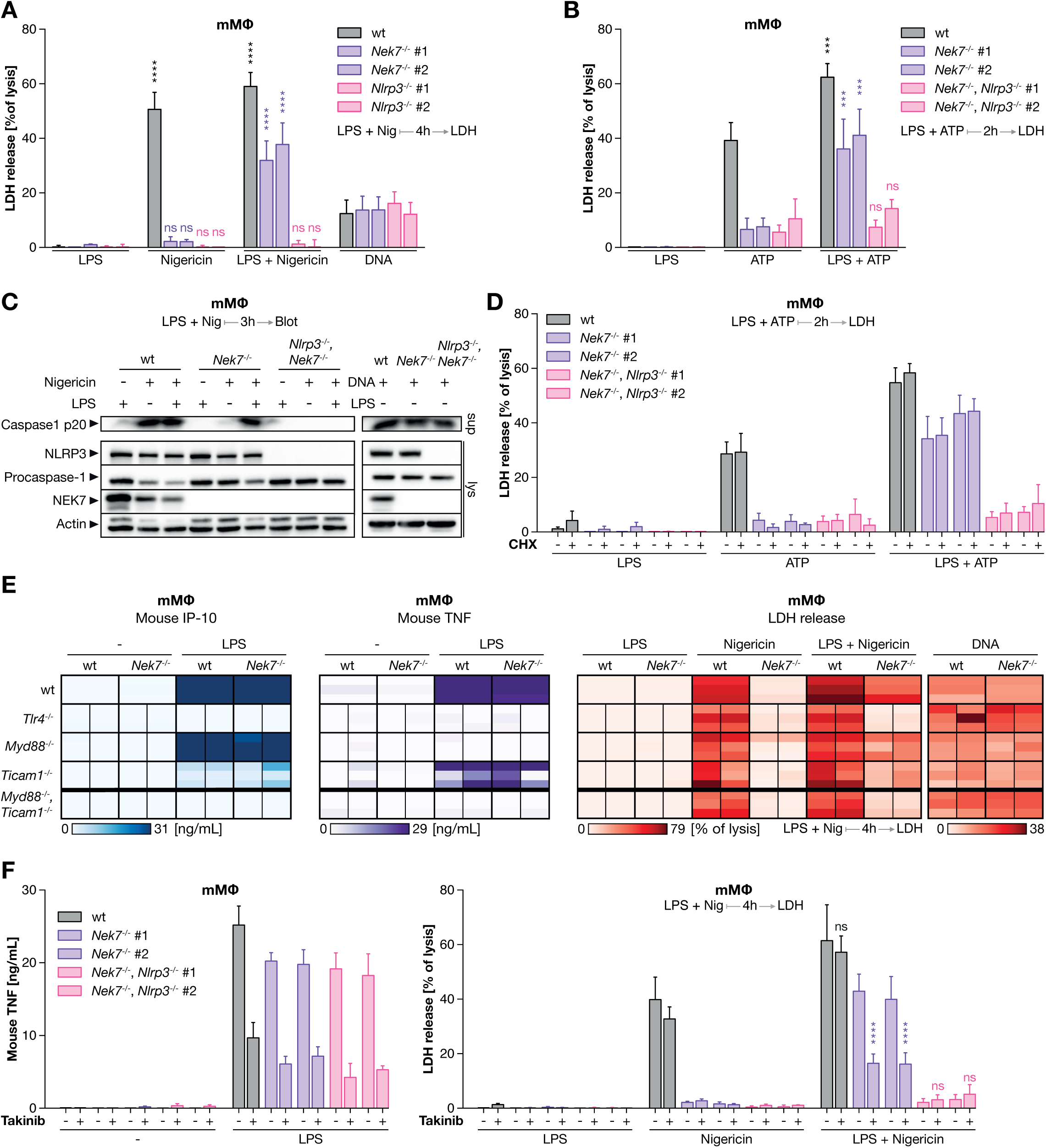
Priming activates TAK1 to bypass NEK7 via a translation-independent mechanism. (A-C) Murine macrophages of the indicated genotypes were stimulated with LPS + Nigericin simultaneously for 4 hr, with DNA for 28 hr or with LPS + ATP for 2 hr. (C) One immunoblot representative of two clones from two independent experiments is shown. (D) mMΦ were pretreated with cycloheximide (CHX) for 30 min and stimulated as in A-C. (E) Two mMΦ clones per genotype were stimulated as indicated. Cytokine and LDH release of two clones (sub-columns) from three independent experiments (sub-rows) are depicted as heatmaps. (F) mMΦ of the indicated genotypes were stimulated as in (A-C) in the presence of Takinib. Bars represent mean + SEM of three independent experiments. **** p < 0.0001, *** p < 0.001, ns p ≥ 0.05. See also Figure S2.

### Human NEK7 can be a cofactor for NLRP3

Turning back to the human innate immune system, we wished to explore whether human myeloid cells activated the NLRP3 inflammasome in the absence of NEK7 due to a species-specific difference in NEK7 or NLRP3 themselves. Immunoprecipitating NLRP3 in THP-1 cells, we found NEK7 to associate with NLRP3 in a constitutive manner, independently of K^+^ efflux (Figure 3A, S3A). Since these results established that NEK7 was in principle capable of interacting with human NLRP3, we wondered whether hNEK7 had the ability to activate hNLRP3. To address this question, we reconstituted the NLRP3 inflammasome in *NEK7*^-/-^ HEK-293T cells with human and murine NLRP3 and NEK7 orthologues (Fig. 3B). These experiments showed that human NEK7 can enhance the activation of both murine and human NLRP3 to a similar extent as murine NEK7 (Fig. 3B, C and S3B, C). These results suggested that although NEK7 is able to participate in hNLRP3 activation, human myeloid cell lines do not rely on NEK7 when forming an NLRP3 inflammasome. Corroborating this, human cells could also activate transgenic mNLRP3 in the absence of NEK7 (Fig. 3D, E). This indicated that it is not a species-specific feature of the NEK7 or NLRP3 molecules themselves that determines whether NLRP3 activation can proceed in the absence of NEK7. Instead, human myeloid cell lines rely by default on a NEK7-independent NLRP3 activation pathway. Based on our observations in murine macrophages that TAK1 activation could bypass NEK7 requirement, we reasoned that BLaER1 cells may employ this pathway by default. Indeed, NLRP3 activation was fully abrogated in TAK1-deficient (*MAP3K7*^-/-^) BLaER1 cells, while NAIP/NLRC4 inflammasome activation remained unaffected (Fig. 3F, G). Importantly TAK1 was not required for transcriptional priming of NLRP3 in BLaER1 cells since NLRP3 expression was fully intact in *MAP3K7*^-/-^ cells (Fig. S3D). Moreover, blocking de novo gene expression using cycloheximide did not impact on NLRP3 inflammasome activation in BLaER1 cells (Fig. S3E). As such, the complete defect of these cells in NLRP3 inflammasome activation suggests that BLaER1 cells are not capable of NEK7-dependent NLRP3 activation. Taken together, these data establish that human NEK7 is able to participate in NLRP3 inflammasome activation but that human myeloid cell lines employ TAK1 to drive post-translational priming of NLRP3 instead.

**Figure 3.**
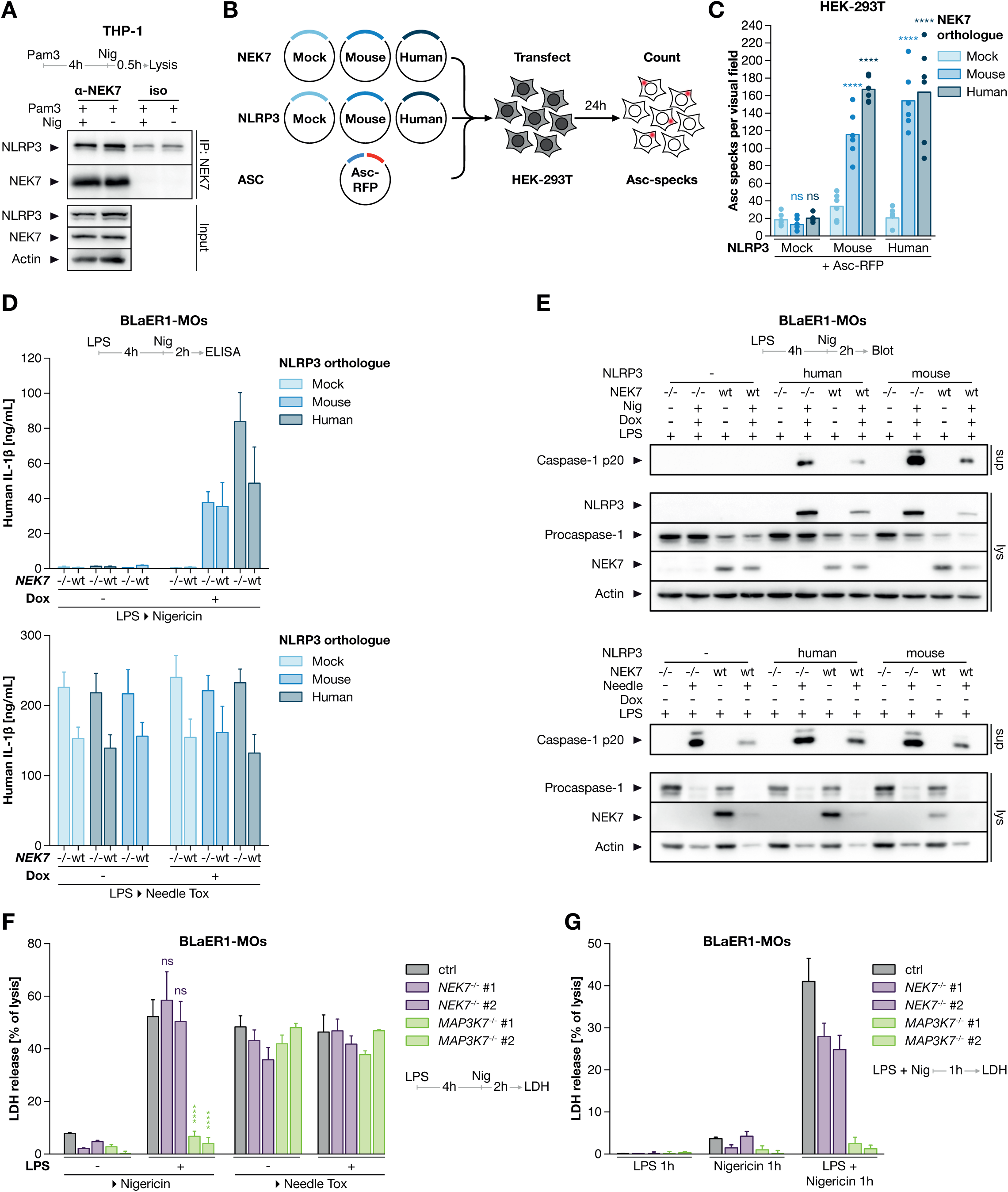
Human NEK7 can be a cofactor for NLRP3. (A) Pam3CSK4-primed THP-1 cells were stimulated with Nigericin for 30 min and lysates were immunoprecipitated with anti-NEK7 or isotype control. One representative immunoblot of three independent experiments is shown. (B, C) *NEK7*^-/-^ HEK293T cells were transiently transfected with the indicated plasmids and ASC-RFP specks were imaged 24 hr after transfection. Dots represent technical replicates from one representative of three independent experiments. (D, E) BLaER1 cells expressing doxycyclin-inducible NLRP3 were stimulated as indicated. Data is depicted as mean + SEM of n=3 (D) or one representative immunoblot of three (E). (F-G) BLaER1 monocytes of the indicated genotypes were primed with LPS for 4 hr and stimulated as indicated for 2 hr (F) or stimulated with LPS + Nigericin for 1 hr (G). Mean + SEM of three independent experiments. **** p < 0.0001, *** p < 0.001, ns p ≥ 0.05. See also Figure S3 and Table S1.

### NEK7 accelerates PTM-primed NLRP3 inflammasome activation in human iPS-Macs

Finally, we wanted to address what role NEK7 has in NLRP3 inflammasome activation in primary human macrophages. To this end we adopted a recently described in vitro differentiation protocol, in which human iPS cells are differentiated into macrophages (hiPS-Macs) (Takata et al., 2017). hiPS-Macs were fully capable of inflammasome activation. Stimulating LPS-primed hiPS-Macs with Nigericin or Needle-Toxin, we observed pyroptosis (LDH release) accompanied by the release of IL-1β and IL-18 (Fig. 4A). Of note, both cytokine- and LDH release in response to Nigericin, but not Needle Tox, were sensitive to treatment with the NLRP3 inhibitor MCC950 (Fig. 4A). To investigate the role of NEK7 in NLRP3 inflammasome activation in hiPS-Macs, we generated *NEK7*^-/-^ iPS cell clones via CRISPR/Cas9 genome editing (Figure 4B). NEK7 deficiency neither impacted on macrophage differentiation nor did it affect NLRP3 expression levels (Fig. 4C). Following differentiation, we stimulated hiPS-Macs with different types of inflammasome stimuli. As readouts for inflammasome activation, we focused on pyroptosis and IL-18 release. IL-1β release could only be studied in cells subjected to prolonged LPS treatment, consistent with the fact that it needs to be induced de novo, similar to other pro-inflammatory cytokines such as IL-6 (Fig. S4A, B). NEK7 dependency was only observed when hiPS-Macs were stimulated with LPS and Nigericin for one hour (Fig. 4D, purple bars), while after four hours of stimulation NLRP3 inflammasome activity was mostly NEK7-independent (Fig. 4D, red bars). Moreover, prolonged LPS (4 hr) treatment followed by Nigericin (2 hr) stimulation was also largely NEK7 independent (Fig. 4D, blue bars). Altogether, these experiments demonstrated that human hiPS-Macs employ NEK7-independent and -dependent modes of NLRP3 activation. Analogous to murine macrophages, concomitant LPS and Nigericin stimulation displays NEK7-dependency at an early time point, while NEK7 is redundant at later time points following NLRP3 activation. After prolonged LPS-treatment, a priming condition typically used for murine macrophages to study IL-1β maturation, a partial NEK7 dependence is observed (Fig. 4E).

**Figure 4.**
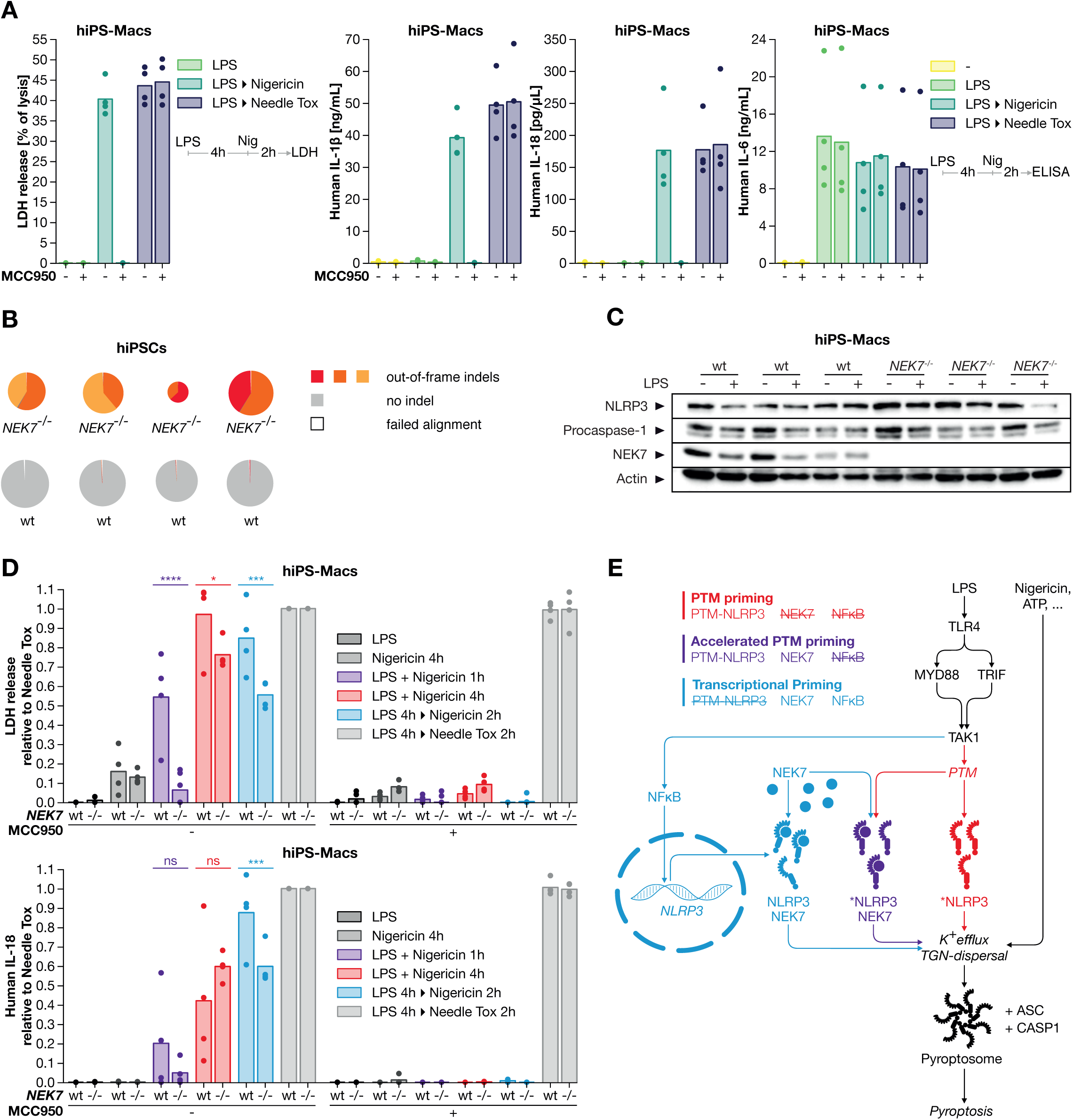
Priming bypasses NEK7 in Human iPS cell derived macrophages. (A) HiPS-Macs were primed with LPS for 4 hr and stimulated as indicated for 2 hr. Cytokine and LDH release of individual clones differentiated in one batch is depicted. (B) Deep sequencing analysis of genome editing in human iPS cells. (C) Immunoblotting of human hiPS-Macs. (D) Wildtype and *NEK7*^-/-^ hiPS-Macs were stimulated as indicated. Dots represent individual clones from one experiment. **** p < 0.0001, *** p < 0.001, * p < 0.05, ns p ≥ 0.05. (E) Model of NEK7-dependent and -independent NLRP3 Priming. PTM: posttranslational modification. See also Figure S4.

## DISCUSSION

It is generally accepted that NLRP3 activation requires two signals. A first signal functions to prime NLRP3 to reach a state that allows it to be activated by a second signal - the actual activation signal. NEK7 has been shown to be an essential cofactor of NLRP3 activation and it has been suggested that NEK7 facilitates inflammasome formation by mediating recognition of the second signal. Studying the role of NEK7 in human myeloid cell lines, we made the unexpected discovery that NLRP3 activation can be fully operational in the absence of NEK7. By genetically dissecting NLRP3 inflammasome signaling we uncovered that these cells employ a NEK7-independent signaling cascade instead that drives TAK1-dependent, post-translational priming of NLRP3. While this TAK1-dependent priming signal is the default pathway by which these human myeloid cell lines engage the NLRP3 inflammasome, murine macrophages predominantly rely on NEK7 for NLRP3 priming. However, they can bypass NEK7 and switch to TAK1-dependent priming under certain stimulatory conditions. This NEK7-independency in human myeloid cells could not be attributed to species-specific constitutions of the NEK7 or NLRP3 molecules themselves. As such, immunoprecipitation and reconstitution experiments showed that human NEK7 interacted with human NLRP3 and that NEK7 was able to facilitate NLRP3 activity. In line with this notion, primary human macrophages also employ NEK7 to activate the NLRP3 inflammasome, yet this requires LPS priming and is only observed at early timepoints, when their default TAK1 post-translational priming mechanism is not yet operational. Indeed, in these cells after prolonged LPS-priming the importance of NEK7-dependent priming declines when the other priming cascade becomes available. Altogether, this study establishes NEK7 as a priming rather than an activation signal. Moreover, in its capacity as a priming factor NEK7 does not constitute an absolute requirement for NLRP3 inflammasome activation. Instead, a priming signal emanating from TAK1 can fully compensate for NEK7. Intriguingly, this signal supersedes NEK7 requirement in human myeloid cell lines and also represents the dominant priming entity in primary human macrophages.

Ever since its first description in 2001, NLRP3 has attracted much attention as a key driver of antimicrobial and sterile inflammation (Gaidt and Hornung, 2018). Nonetheless, despite being in the focus for almost two decades the molecular mechanism of NLRP3 activation has remained obscure. The two-step model of inflammasome activation predates the discovery of NLRP3 and inflammasomes altogether, originating from the notion that both a pro-inflammatory and a cell-death inducing signal are required to release mature IL-1β from murine bone marrow derived macrophages (Hogquist et al., 1991). In retrospect, these early studies had assessed NLRP3 inflammasome activation employing a K^+^ efflux inducing trigger. Subsequent studies revealed that the pro-inflammatory signal indeed serves two independent functions in the context of NLRP3 inflammasome activation. While this signal is critically required to induce pro-IL-1β expression, it is also necessary to render NLRP3 activatable in the first place. This became apparent when studying the maturation of caspase-1, the expression of which is independent of a proinflammatory signal, as a proxy of NLRP3 inflammasome activation. Here it was revealed that unprimed macrophages did not mature caspase-1 upon K^+^ efflux inducing stimuli (Kahlenberg et al., 2005; Mariathasan et al., 2004), but that additional priming by a proinflammatory signal was required to facilitate this step. Of note, this unique requirement of NLRP3 priming by a pro-inflammatory signal (referred to as signal 1 or priming in this manuscript) must not be confused with the signal that induces pro-IL-1β expression. Indeed, while both signals can be provided by the same PRR, they can also be separated, and the pro-IL-1β inducing stimulus is not necessary for NLRP3 inflammasome activation.

Although the two-step activation model constitutes an important conceptual framework for NLRP3 activation, it has proven to be an enormous conundrum because it is not trivial to allocate signaling events upstream of NLRP3 to either priming or activation. The fact that several NLRP3 priming pathways have been described (Groslambert and Py, 2018) is likely attributable to the fact that stimulus, cell type, species and also temporal aspects play an important role in this context. We conceptualize that priming serves the function to increase the pool of NLRP3 molecules that are able to respond to an activating stimulus, or to lower the activation threshold of individual NLRP3 molecules. In this regard, we would interpret the existence of multiple redundant NLRP3 priming pathways as the possibility to integrate diverse pro-inflammatory inputs to achieve this activatable state. In fact, we consider this pleiotropy to be a key trait of priming - but not activation pathways. Based on the redundancy between TAK1 and NEK7 in facilitating NLRP3 inflammasome activation, we conclude that NEK7 serves as a priming factor of the NLRP3 inflammasome. In line with this notion, recent structural evidence suggests that NEK7 functions to stabilize individual NLRP3 molecules in a semi-open, but not fully active conformation (Sharif et al., 2019). Conceivably, priming-related post-translational modifications might function in a similar *modus operandi* and are thus redundant with NEK7, as observed in mouse macrophages, or synergistic, as observed in primary human macrophages.

NEK7 holds a unique position amongst NLRP3 priming pathways in that it is constitutively expressed and apparently uncoupled from upstream signals in its pro-inflammatory capacity. In this regard, it has been suggested that NEK7 is employed for NLRP3 activation in order to avoid inflammasome formation during mitosis, when NEK7 is not available (Shi et al., 2016). Indeed, it has been speculated that the cellular perturbation immediately upstream of NLRP3 commonly occurs during mitosis and thus the dependency on NEK7 prevents inadvertent inflammasome activation during cell division (Sharif et al., 2019). However, the here-uncovered redundancy of NEK7 priming with other cell cycle-independent priming pathways (e.g. TAK1) advocate against a specific de-coupling of NLRP3 inflammasome activation and proliferation. This is also in line with the fact that many NLRP3 inflammasome-competent cells of the innate immune system are indeed postmitotic. As such, despite detailed mechanistic insight into how NEK7 can accelerate NLRP3 inflammasome activation, the physiological role of NEK7 remains to be determined.

Our data suggest that TAK1-dependent signaling constitutes the predominant NLRP3 priming modality in human myeloid cells. Intriguingly, a previous study exploring kinase inhibitors in the context of inflammasome signaling had also identified TAK1 kinase activity as an important facilitator of NLRP3 inflammasome activation (Gong et al., 2010). While these data would argue for TAK1 kinase activity as a potential drug target in the NLRP3 inflammasome pathway, it has to be noted that prolonged inhibition of TAK1 can result in the induction of lytic cell death in the context of pro-inflammatory signals (Malireddi et al., 2018). This, in turn, can lead to secondary activation of the NLRP3 inflammasome due to the perturbation of membrane integrity and associated K^+^ efflux. Consequently, directly targeting NLRP3, or the TAK1-dependent conversion of NLRP3 into an activatable state, would constitute a safer pharmacological approach towards inhibition of NLRP3.

## MATERIALS & METHODS

### BLaER1 cell culture

BLaER1 cells (female) were cultivated in RPMI supplemented with 10 % FCS, 1 mM Pyruvate, 100 U/mL Penicillin and 100 µg/mL Streptomycin at 37 °C and 5 % CO_2_. BLaER1 cells were differentiated in Medium containing 10 ng/mL hrIL-3, 10 ng/mL hrCSF-1 (M-CSF) and 100nM β-Estradiol for 5-6 days. In the course of these studies, we serendipitously identified that BLaER1 cells express transcripts of SMRV (Squirrel monkey retrovirus) and subsequent experiments confirmed that BLaER1 cells harbor the SMRV proviral genome. Testing early passages of BLaER1 cells by Dr. Thomas Graf (personal communication) confirmed that the parental BLaER1 cell line (Rapino et al., 2013) is positive for SMRV. Of note, extensive characterization of BLaER1 monocytes in comparison to other human myeloid cells has not provided any indication that SMRV positivity would impact on the functionality of these cells as myeloid cells. Samples of other cell lines used in this work were confirmed to be free of SMRV by PCR.

### THP-1 cell culture

THP-1 cells (male) were obtained from ATCC and cultivated in RPMI supplemented with 10 % FCS, 1 mM Pyruvate, 100 U/mL Penicillin and 100 µg/mL Streptomycin at 37 °C and 5 % CO_2_. THP-1 cells were differentiated by adding 100 ng/ml PMA to the medium for 18 hr, rinsed off with ice-cold PBS and replated for experiments.

### hiPSC, hiPS-Macs cell culture

Human induced pluripotent stem cells (hiPSCs) were kindly provided by Adam O’Neill and Magdalena Götz (Camargo Ortega et al., 2019). hiPSCs were cultivated on Geltrex-coated plates in complete mTeSR1 Medium at 37 °C and 5 % CO_2_ and detached for passaging using 1.5mL Accutase for 5 minutes at 37 °C after a PBS wash. After passaging, cells were cultivated in the presence of 5 µM ROCK-Inhibitor overnight.

### Murine Macrophage cell culture

Murine Macrophages were cultivated in DMEM supplemented with 10 % FCS, 1 mM Pyruvate 100 U/mL Penicillin and 100 µg/mL Streptomycin at 37 °C and 5 % CO_2_. mMΦ were detached for passaging with 0.05 % Trypsin at 37° for 15 minutes after one PBS wash and then rinsed off with DMEM.

### HEK-293T cell culture

HEK-293T cells were cultivated in DMEM with 10 % FCS, 1 mM Pyruvate, 100 U/mL Penicillin and 100 µg/mL Streptomycin at 37 °C and 5 % CO_2_. For passaging, cells were washed with PBS once and then incubated with 0.05 % Trypsin at 37° for 5 minutes. Cells were then rinsed off with DMEM.

### ASC pyroptosome imaging

ASC pyroptosomes in transiently transfected HEK-293T cells were imaged 24 hr after Transfection on a Leica Hi8 epifluorescence microscope using 10x magnification. Specks were quantified with CellProfiler (Carpenter et al., 2006).

### Differentiation of hiPSCs into hiPS-Macs

Differentiation into iPS-Macs was achieved as described previously (Takata et al., 2017). Briefly, 150.000 hiPSC were plated into a one well of a Geltrex-coated 6well plate and differentiated in StemPro base medium with StemPro Supplement, 200 µg/mL Human Transferrin, 2 mM Glutamine, 0.45 mM MTG and 0.5 mM Ascorbic Acid (= StemPro medium, ascorbic acid was added just before use) by stimulation with 50 ng/mL VEGF, 5 ng/mL BMP-4 and 2 mM CHIR99021 at 5 % oxygen for two days, followed by two days of stimulation with 50 ng/mL VEGF, 5 ng/mL BMP-4 and 20 ng/mL FGF2. From day four, StemPro medium was supplemented with 15 ng/mL VEGF and 5 ng/mL FGF2. Starting at day six, 10 ng/mL VEGF, 10 ng/mL FGF2, 50 ng/mL SCF, 30 ng/mL DKK-1, 10 ng/mL IL-6 and 20 ng/mL IL-3 were added to StemPro medium until day ten. From day eight, cells were cultivated under normoxic conditions. From day twelve, 10 ng/mL FGF2, 50 ng/mL SCF, 10 ng/mL IL-6 and 20 ng/mL IL-3 were added to StemPro medium. Starting at day sixteen, cells were cultivated in 75 % IMDM with 25 % F12 supplement, N2 supplement, B-27 supplement, 0.05 % BSA and 100 U/mL Penicillin and 100 µg/mL Streptomycin (= SF-Diff Medium) supplemented with 50 ng/mL rhCSF-1 (M-CSF) at least until day 28. Culture medium was exchanged as necessary, but at least every two days. After differentiation, hiPS-Macs were carefully harvested from the supernatant, spun down and replated in RPMI medium with 10 % FCS, 1 mM Pyruvate, 100 U/mL Penicillin and 100 µg/mL Streptomycin for experiments.

### Immunoblotting

Cells were lysed at approximately 5 Mio/mL in 1x Lämmli Buffer and boiled for 5 minutes at 95 °C. For precipitation of total protein from supernatants, stimulations were done in medium containing 3% FCS. Precipitation of total protein from supernatants was achieved by combining 700 µL of supernatant with 700 µL MeOH and 150 µL of CHCl_3_. Samples were spun down at 20.000g for 20 minutes, and the upper phase was discarded. Again, 700 µL MeOH were added and samples were centrifuged at 20.000 g for 20 minutes. The pellet was then dried and resuspended in 100 µL 1x Lämmli buffer and boiled at 95 °C for 5 minutes. Samples were run on 12 % SDS-PAGE gels at 150 V for 85 minutes and were subsequently transferred onto a nitrocellulose membrane at 100 V for 75 minutes at 4 °C. Membranes were then blocked in 5 % milk for 1 hr at room temperature. Primary and secondary antibodies were diluted in 1-5 % milk.

### ELISA and LDH assay

LDH assays were done on supernatants immediately after experiments. Results are presented relative to a lysis control from the same experiment with the values of unstimulated controls subtracted as background. ELISAs were done according to manufacturer’s instructions on supernatants stored at −20 °C.

### Stimulation

NLRP3 was primed as indicated with 1 µg/mL Pam3CSK4 or 200 ng/mL LPS. NLRP3 was activated with 5 mM ATP for 2 hr or Nigericin at 6.5 µM or 10 µM as indicated for up to 4 hr. To activate the AIM2 inflammasome 400ng HT-DNA were transfected into a 96-well with 0.5 µL Lipofectamine in 50 µL OptiMEM by incubating OptiMEM and Lipofectamine for 5 minutes followed by 20 minutes of incubation of the Lipofectamine-DNA mix in OptiMEM and dropwise addition of the mix to the cells. The NAIP/NLRC4 inflammasome was activated with 0.25 µg/mL protective antigen (pA) and 0.025 µg/mL needle tox (LF-YscF) for 2 hr.

### Inhibition of translation

For murine macrophages, cycloheximide (CHX) was added to the medium 30 min before stimulation to a final concentration of 10 µg/mL. For BLaER1 cells, CHX was added to the medium simultaneously with LPS at the indicated concentrations in the range of 1-10 µg/mL.

### Doxycyclin-inducible gene expression

In BLaER1 cells transduced with pLI-Puro derived vectors, gene expression was induced by adding Medium to a final concentration of 1 µg/mL doxycyclin for the last 24 hr of differentiation.

### Pharmacological Inhibition of TAK1 and NLRP3

Takinib was added to the medium to a final 50 µM concentration 1 hr before stimulation. MCC950 was added to the medium 1 hr before priming to a final concentration of 10 µM.

### Inhibition of K^+^-efflux

Potassium chloride was added to the medium together with the priming stimulus to the indicated final concentrations. The overall osmolarity of the medium was kept constant over all conditions.

### Transient Transfection of HEK-293T cells

HEK-293T cells were transiently transfected with 400ng Plasmid DNA in 50 µL OptiMEM with 1 µL GeneJuice by incubating GeneJuice with OptiMEM for 5 minutes followed by 15 minutes of incubation of the DNA-GeneJuice mix in OptiMEM. DNA concentrations were kept constant across all conditions using pBluescript as stuffer DNA.

### Plasmid DNA purification

Plasmid DNA was purified from *E.Coli* DH5α using a Thermo HiPure Maxiprep Kit according to manufacturer’s instructions.

### Preparation of Lentiviral Particles

Lentiviral particles were prepared according to (Kutner et al., 2009). Briefly, HEK-293T cells were transfected with 20µg transfer plasmid, 15µg pCMVΔ8.91 packaging plasmid and 6µg pMD2.G VSV-G pseudotyping plasmid dish by diluting the plasmids in 1 mL 1x HBS, adding 50 µL 2.5 M Calcium chloride and gently pipetting the mix onto a 10cm dish with approximately 6 Mio. HEK-293T cells in fresh medium. After 8 hr the medium was exchanged. Supernatants were harvested 48 hr later, spun down and filtered before being used to transduce target cells. Successfully transduced cells were selected with 2.5 - 5 µg/mL Puromycin or 10 µg/mL Blasticidin S for 48 hr, or FACSorted for Fluorescence markers.

### Genome Editing and Overexpression

sgRNA oligos were designed using CHOPCHOP (Labun et al., 2019) and cloned into expression Plasmids as described previously (Sanjana et al., 2014; Schmid-Burgk et al., 2014). BLaER1 cells were electroporated in OptiMEM with 5 µg of plasmids driving expression of Cas9 and an sgRNA on a BioRad GenePulser XCell as described previously (Schmidt et al., 2015). THP-1 cells and murine macrophages were transduced with lentiviral particles driving expression of Cas9 (Lenti-Cas9-Blast (Sanjana et al., 2014)) or an sgRNA (LentiGUIDE-Puro (Sanjana et al., 2014)). HEK-293T cells were transiently transfected with plasmids driving expression of Cas9 or an sgRNA. LentiGuide-Puro (Addgene plasmid #52963) and LentiCas9-Blast (Addgene plasmid #52962) were gifts from Feng Zhang.

hiPS cells conditioned to grow as single clones were electroporated with Cas9-crRNA-trRNA complexes (RNPs) targeting *NEK7*. Grown single clones were duplicated, lysed and out-of-frame editing in *NEK7* was analyzed via deep sequencing as described previously (Schmid-Burgk et al., 2014). Several *NEK7^-/-^* and *NEK7^+/+^* clones were expanded and used for experiments.

## STATISTICAL ANALYSIS

Numbers of independent replicates (n) are reported in the respective figure legends. p-values were calculated based on two-way ANOVAs followed by Sidak’s multiple comparisons test for groups containing two elements, or Tukey’s test for larger groups. All statistical analyses were done using GraphPad Prism 7. * p < 0.05, ** p < 0.01, *** p < 0.001, **** p < 0.0001, ns p ≥ 0.05.

## RESOURCES

| REAGENT or RESOURCE | SOURCE | IDENTIFIER |
| --- | --- | --- |
| **Antibodies** |  |  |
| anti-Caspase-1 (p20) (human), mAb (Bally-1) | AdipoGen, San Diego, CA | Cat# AG-20B-0048-C100 |
| anti-Caspase-1 (p20) (mouse), mAb (Casper-1) | AdipoGen | Cat# AG-20B-0042-C100 |
| anti-NEK7 | Abcam, Cambridge, UK | Cat# ab133514 |
| anti-NLRP3/NALP3, mAb (Cryo-2) | AdipoGen | Cat# AG-20B-0014-C100 |
| anti-Human IL-1 beta /IL-1F2 | R&D Systems Inc, Minneapolis, MN | Cat# AF-201-NA |
| **Chemicals, Peptides, and Recombinant Proteins** |  |  |
| 1-Thioglycerol (MTG) | Sigma-Aldrich, St. Louis, MO | Cat# M6145 |
| Accutase | Stemcell Technologies, Vancouver, Canada | Cat# 07920 |
| Adenosine 5'-triphosphate disodium salt hydrate | Sigma-Aldrich | Cat# A6419 |
| Ascorbic Acid | Sigma-Aldrich | Cat# A4544-100G |
| B-27 supplement | Thermo Fisher Scientific, Waltham, MA | Cat# 17504-001 |
| Blasticidin S HCl (10 mg/mL) | Thermo Fisher Scientific | Cat# A1113903 |
| BSA | GE Healthcare, Chicago, IL | Cat# SH30574.01 |
| CHIR99021 | Miltenyi Biotec, Bergisch Gladbach, Germany | Cat# 130-103-926 |
| Cycloheximide | Carl Roth, Karlsruhe, Germany | Cat# 8682.1 |
| Doxycycline hyclate | Sigma-Aldrich | Cat# D9891-1G |
| Geltrex | Thermo Fisher Scientific | Cat# A1413302 |
| GeneJuice | Merck, Darmstadt, Germany | Cat# 70967-3 |
| Glutamic Acid | Thermo Fisher Scientific | Cat# 25030024 |
| Ham's F12 nutrient mix | Thermo Fisher Scientific | Cat# 21765029 |
| Herring Testis(HT)-DNA sodium salt | Sigma Aldrich | Cat# D6898 |
| Human CSF-1 (M-CSF) (iPSC differentiation) | R&D Systems | Cat# 216-MC-005 |
| Human Transferrin | Roche, Basel, Switzerland | Cat# 10-652-202-001 |
| IMDM with GlutaMAX | Thermo Fisher Scientific | Cat# 31980022 |
| LF-YscF | This Study | N/A |
| Lipofectamine 2000 Transfection Reagent | Thermo Fisher Scientific | Cat# 11668019 |
| LPS-EB Ultrapure | Invivogen, San Diego, CA | Cat# tlrl-3pelps |
| MCC950 | Sigma-Aldrich | Cat# PZ0280 |
| mTeSR1 | Stemcell Technologies | Cat# 85850 |
| N-2 Supplement | Thermo Fisher Scientific | Cat# 17502048 |
| Nigericin sodium salt | Sigma-Aldrich | Cat# N7143 |
| Pam3CSK4 | Invivogen | Cat# tlrl-pms |
| Phorbol 12-myristate 13-acetate | ENZO Life Sciences, Farmingdale, NY | Cat# BML-PE160-Cat# 0005 |
| Protective antigen (pA) | Biotrend, Cologne, Germany | Cat# LL-171E |
| Puromycin Dihydrochlorid | Carl Roth | Cat# 0240.4 |
| Recombinant Human BMP-4 | R&D Systems | Cat# 314-BP-010 |
| Recombinant Human CSF-1 (M-CSF) (BlaER1 differentiation) | Recombinantly produced | N/A |
| Recombinant Human DKK-1 | R&D Systems | Cat# 5439-DK-010 |
| Recombinant Human FGF2 | R&D Systems | Cat# 233-FB-025 |
| Recombinant Human IL-3 | R&D Systems | Cat# 203-IL-010 |
| Recombinant Human IL-3 (BLaER1 differentiation) | Recombinantly produced | N/A |
| Recombinant Human IL-6 | R&D Systems | Cat# 206-IL-010 |
| Recombinant Human SCF | R&D Systems | Cat# 255-SC-010 |
| Recombinant Human VEGF | R&D Systems | Cat# 293-VE-010 |
| ROCK Inhibitor Y-27632 | Stemcell Technologies | Cat# 72302 |
| Stempro-24 SFM | Thermo Fisher Scientific | Cat# 10639-011 |
| Takinib | Selleck Chemicals, Houston, TX | Cat# S8663 |
| β-Estradiol | Sigma-Aldrich | Cat# E8875 |
| Critical Commercial Assays |  |  |
| Human IL-1β ELISA Set II | BD Biosciences, San José, CA | Cat# 557953 |
| Human Total IL-18 DuoSet ELISA | R&D Systems | Cat# DY318-05 |
| MiSeq Reagent Kit v2, 300 Cycles | Illumina, San Diego, CA | Cat# MS-102-2002 |
| Mouse CXCL10/IP-10/CRG-2 DuoSet ELISA | R&D Systems | Cat# DY466 |
| Mouse TNF (Mono/Mono) ELISA Set II | BD Biosciences | Cat# 558534 |
| OptEIA Human IL-6 ELISA Set | BD Biosciences | Cat# 555220 |
| OptEIA Mouse IL-1β Elisa Set | BD Biosciences | Cat# 559603 |
| Pierce LDH Cytotoxicity Assay Kit | Thermo Fisher Scientific | Cat# 88954 |
| Experimental Models: Cell Lines |  |  |
| BLaER1 | (Rapino et al., 2013) | N/A |
| HEK-293T | (Cavlar et al., 2013) | N/A |
| iPSC | (Camargo Ortega et al., 2019) | N/A |
| Mouse Macrophages, *Nlrp3*, *Asc-CFP*, *Cas9*-expressing | (Franklin et al., 2014) | N/A |
| THP-1 | ATCC, Manassas, VA | Cat# TIB-202 |
| Oligonucleotides |  |  |
| *Refer to supplementary Table 2 for sgRNA target sites* |  |  |
| Recombinant DNA |  |  |
| LentiCas9-Blast | (Sanjana et al., 2014) | N/A |
| LentiGUIDE-Puro | (Sanjana et al., 2014) | N/A |
| pBabe-U6-sgRNA-Cas9 | (Schmidt et al., 2015) | N/A |
| pBlast-hNEK7 | This study | N/A |
| pBlast-mNek7 | This study | N/A |
| pLIX-hNLRP3 | This study | N/A |
| pLIX-mNlrp3 | This study | N/A |
| pLK0.1-sgRNA-CMV-GFP | (Schmid-Burgk et al., 2014) | N/A |
| pRP-Asc-RFP | This study | N/A |
| pRZ-CMV-Cas9 | (Schmidt et al., 2015) | N/A |
| Software and Algorithms |  |  |
| CellProfiler 3.1.5 | (Carpenter et al., 2006) | https://cellprofiler.org |
| CHOPCHOP | (Labun et al., 2019) | https://chopchop.cbu.uib.no |
| Outknocker | (Schmid-Burgk et al., 2014) | http://www.outknocker.org |
| Prism 7.0 | GraphPad, San Diego, CA | https://www.graphpad.com/scientific-software/prism/ |

## AUHTOR CONTRIBUTIONS

Conceptualization N.A.S., M.M.G., F.O., J.L.S.-B. and V.H.; Formal Analysis, Software, Visualization N.A.S.; Investigation N.A.S., M.M.G., I.S., C.A.S., D.C., A.L.F., D.N., F.P. and J.L.S.-B.; Resources V.H.; Writing N.A.S., M.M.G. and V.H. with input from all authors; Funding Acquisition V.H.; Supervision V.H.

## ACKNOWLEDGEMENTS

We kindly thank Andreas Wegerer (Gene Center, LMU) for great technical support, the BioSysM FACS Core Facility (Gene Center, LMU) for cell sorting, Andreas Pichlmair (Technical University of Munich, Munich) for providing the pLIX plasmid, Adam O’Neill and Magdalena Götz (Department of Physiological Genomics, LMU) for providing us with the hiPSCs and helping us set up experiments with these cells, Russell Vance (UC Berkeley, USA) for providing us with the Needle Tox expression plasmid and Manuela Moldt and Karl-Peter Hopfner (Gene Center, LMU) for help producing the Needle Tox protein. This work was supported by grants from the ERC (ERC-2014-CoG – 647858 GENESIS) and funded by the Deutsche Forschungsgemeinschaft (DFG, German Research Foundation) – project number 404446805 – TRR 237 to VH.

## SUPPLEMENTAL FIGURE LEGENDS

**Figure S1 related to Figure 1.**
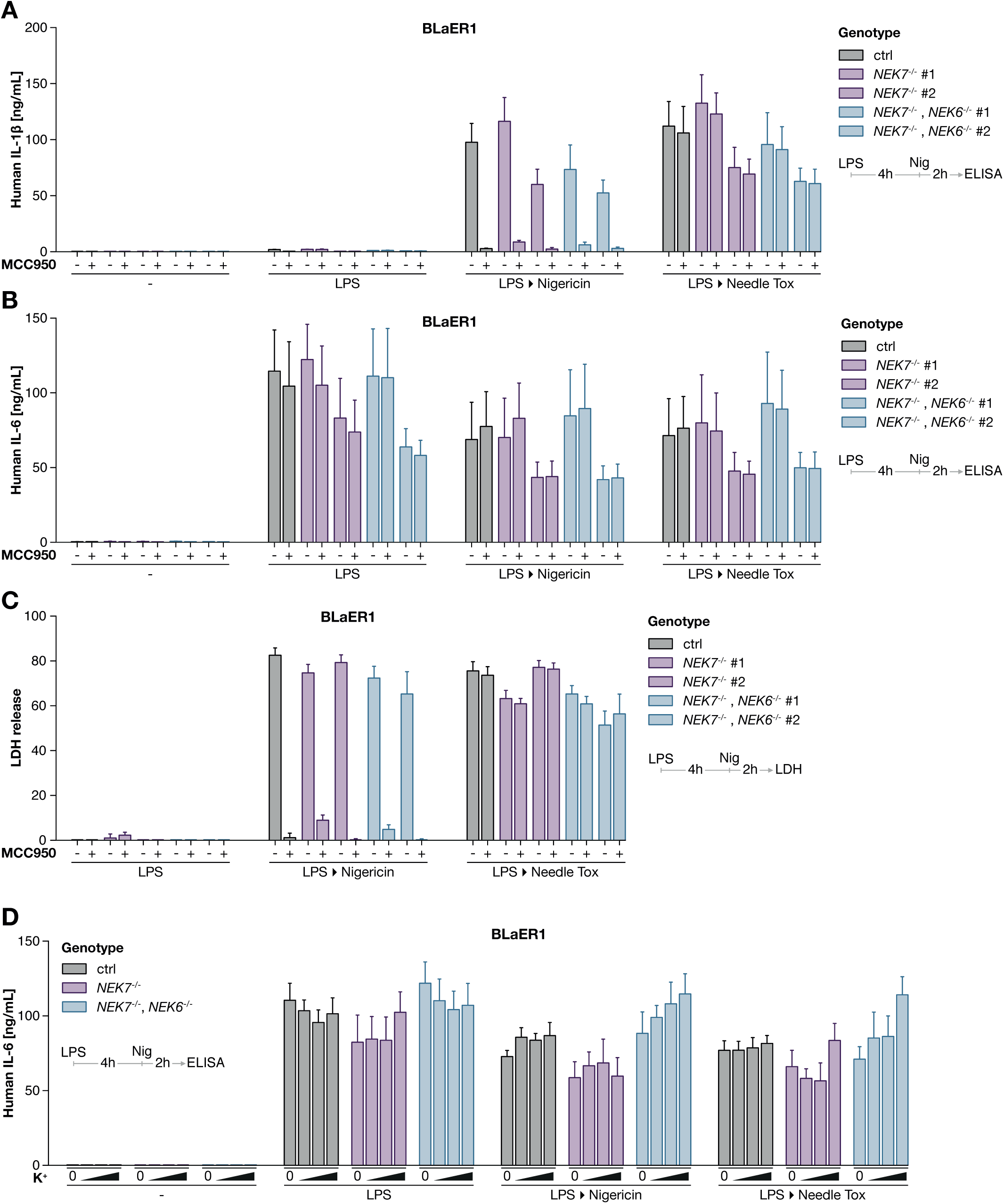
Human myeloid cell lines activate NLRP3 independently of NEK7. (A - C) BLaER1 monocytes were stimulated as in A in the presence of 10 µM MCC950. IL-1β, IL-6 and LDH release are depicted as mean + SEM of three independent experiment. (D) BLaER1 monocytes of the indicated genotypes were primed with 200 ng/mL LPS for 4 hr and subsequently stimulated with 6.5 µM Nigericin or 0.25 µg/mL pA and 0.025 µg/mL Needle Tox (LF-YscF) in the presence of up to 60 mM Potassium Chloride. IL-6 release is depicted as mean + SEM of three independent experiments.

**Figure S2 related to Figure 2.**
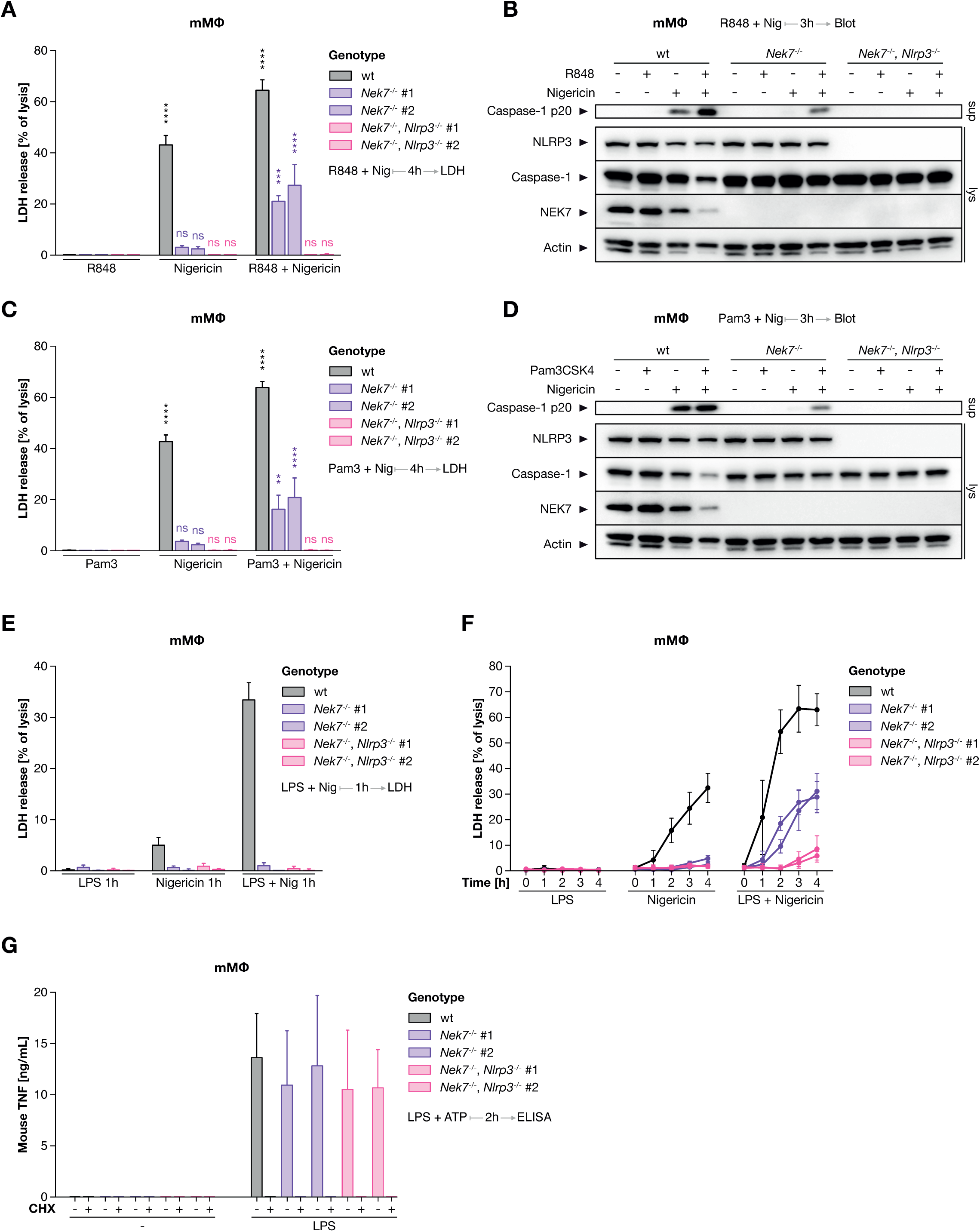
Priming activates TAK1 to bypass NEK7 via a translation-independent mechanism. (A) Murine Macrophages of the indicated genotypes were treated with R848 + Nigericin simultaneously for 4 hr. (B) Murine Macrophages of the indicated genotypes were treated with R848 + Nigericin simultaneously for 3 hr. (C) Murine Macrophages of the indicated genotypes were treated with Pam3CSK4 (Pam3) + Nigericin simultaneously for 4 hr. (D) Murine Macrophages of the indicated genotypes were treated with Pam3CSK4 + Nigericin simultaneously for 3 hr. (E) Murine Macrophages of the indicated genotypes were treated with LPS + Nigericin simultaneously for 1 hr. (F) Murine Macrophages of the indicated genotypes were stimulated as indicated for up to 4 hr. (G) Murine macrophages of the indicated genotypes were pretreated with cycloheximide (CHX) and stimulated with LPS for 2 hr. Release of TNF and LDH is depicted as the mean of three independent experiments ± SEM. Western Blots represent one of two clones from one of three independent experiments. **** p < 0.0001, *** p < 0.001, ** p < 0.01.

**Figure S3 related to Figure 3.**
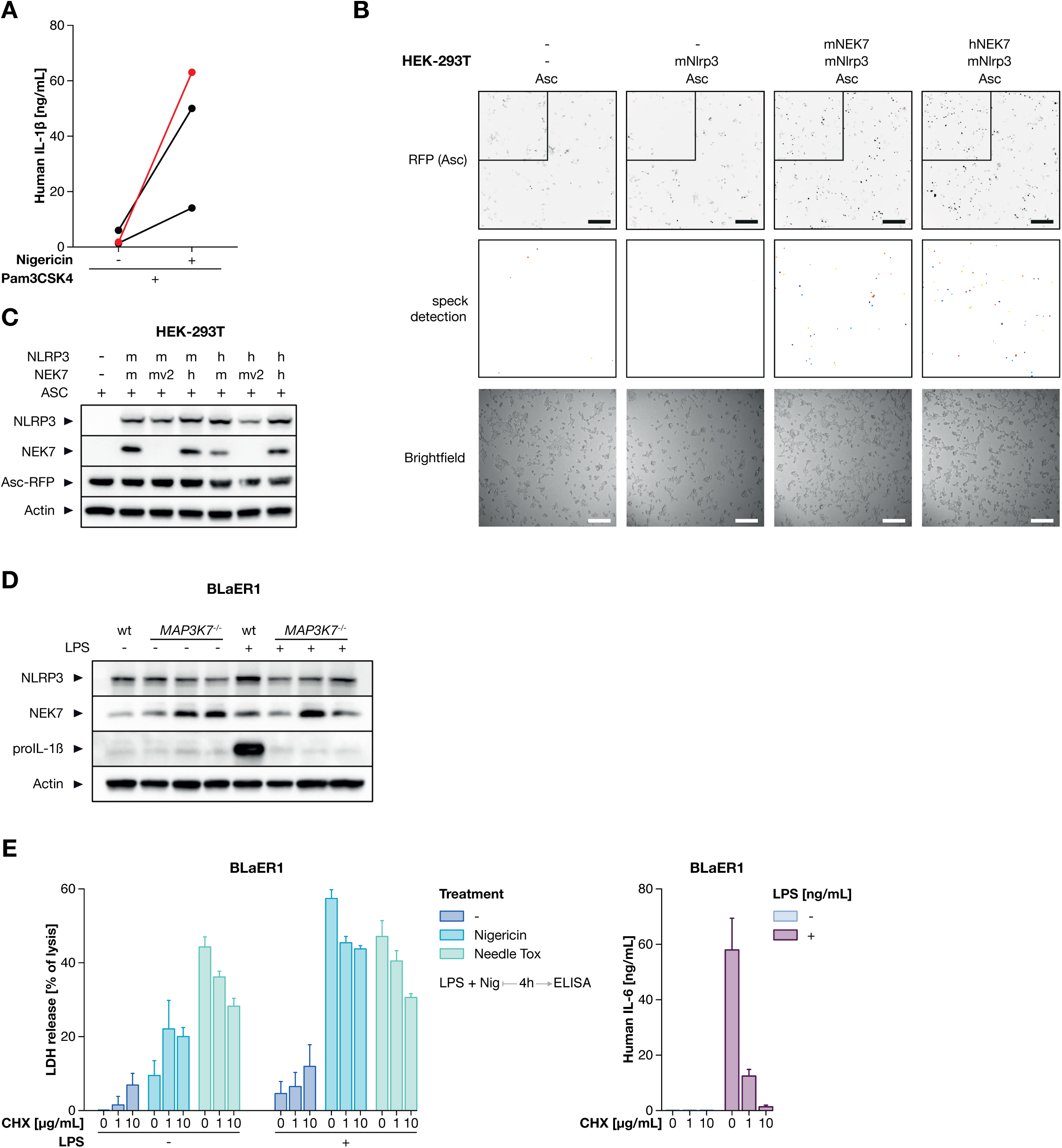
Human NEK7 can be a cofactor for NLRP3. (A) Dots represent IL-1β release from replicates of the immunoprecipitation in Figure 3A. Data in red correspond to the replicate shown in Figure 3A. (B) Sample images corresponding to the experiment shown in Figure 3C. Colors of RFP fluorescence images were inverted for visual clarity, scalebars represent 300 µm. “speck detection” shows automatically detected specks in the area outlined in “RFP”. (C) The expression of NEK7, NLRP3 and ASC-RFP was confirmed via immunoblotting 24 hr after transient transfection of HEK-293T cells. Immunoblot is one representative of two independent experiments. (D) TAK1 deficient BLaER1 Monocytes were treated with LPS for 6 hr before lysates were immunoblotted. One representative of three independent experiments is shown. (E) BLaER1 monocytes were primed with LPS and treated with Nigericin or protective antigen and needle tox in the presence of the indicated concentrations of CHX. LDH and IL-6 release are depicted as mean + SEM of three independent experiments.

**Figure S4 related to Figure 4.**
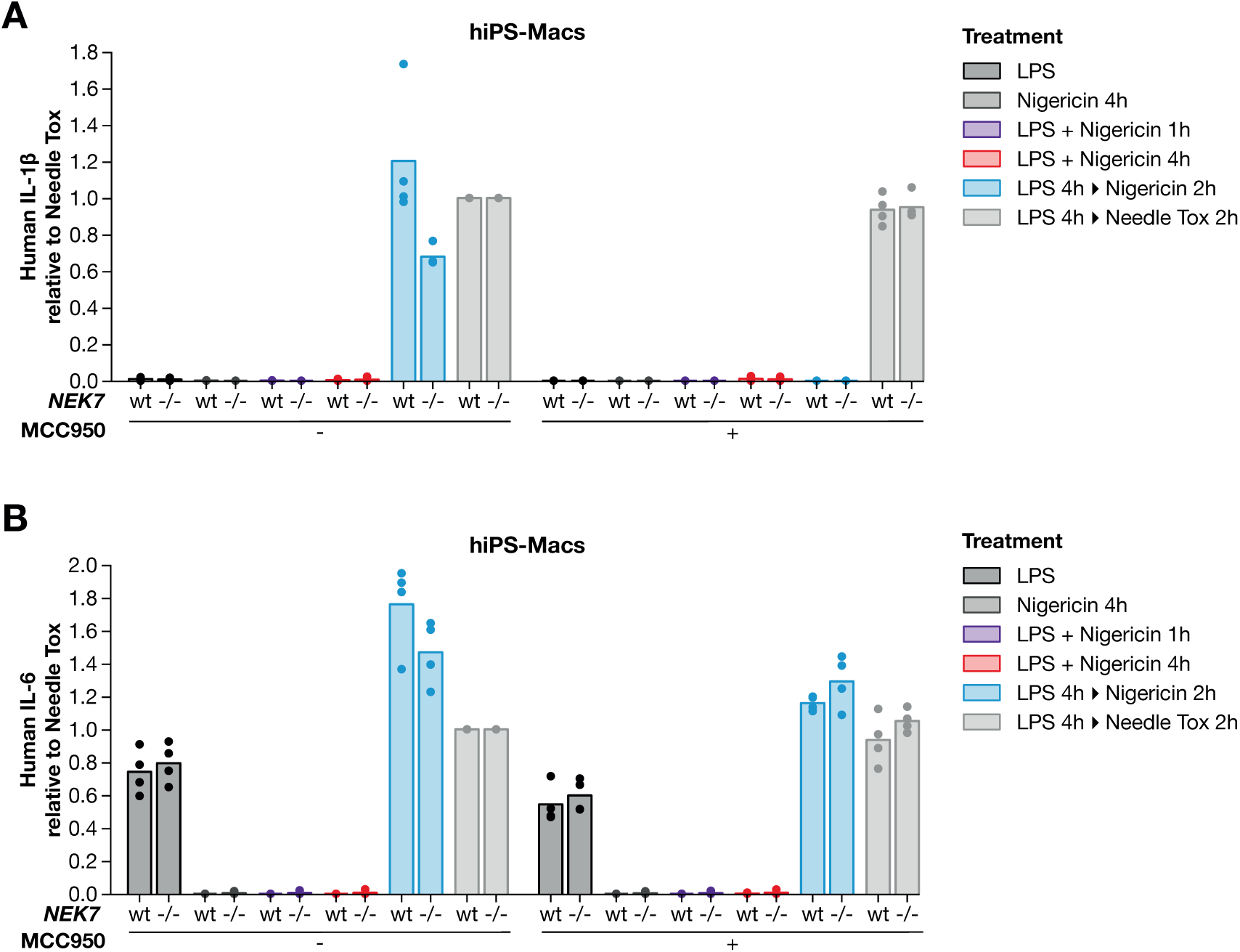
Priming bypasses NEK7 in Human iPS cell derived macrophages. (A, B) wt and *NEK7^-/-^* hiPS-Macs were stimulated as indicated. Release of IL-1β (A) and IL-6 (B) is depicted as mean + SEM of three independent experiments.

